# Colonic metabolomic and transcriptomic alterations in a mouse model of metabolic syndrome

**DOI:** 10.64898/2026.04.02.716131

**Authors:** Jaclyn A. Rivas, David P. Scieszka, Eduardo Peralta-Herrera, Crystal Madera Enriquez, Seth D. Merkley, Audrey L. Nava, Rama R. Gullapalli, Eliseo F. Castillo

**Affiliations:** Division of Gastroenterology and Hepatology, Department of Internal Medicine, University of New Mexico Health Sciences, Albuquerque, NM; Department of Pathology, University of New Mexico Health Sciences, Albuquerque, NM; Comprehensive Cancer Center, University of New Mexico, Albuquerque, NM

**Keywords:** Metabolic Syndrome, colon, metabolomics, transcriptomics, gut microbiotaKeywords: Metabolic Syndrome, colon, metabolomics, transcriptomics, gut microbiota

## Abstract

Metabolic syndrome (MetS), characterized by abdominal obesity, insulin resistance, dyslipidemia, and hypertension, affects a substantial proportion of the global population and increases the risk for cardiovascular disease, diabetes, and metabolic dysfunction-associated steatotic liver disease (MASLD). Despite its prevalence, there are currently no effective pharmacological therapies targeting MetS, highlighting the need to identify novel etiological mechanisms, particularly within the gastrointestinal (GI) tract. Using a mouse model of MetS and healthy lean controls, we assessed the colonic microenvironment through metabolomic, transcriptomic, and microbiome analyses. Colonic organoids were cultured to further explore epithelial alterations. Additionally, human MetS fecal metabolomics data were cross-compared with the mouse model to validate translational relevance. MetS mice exhibited upregulation of colonic anabolic pathways, including glycolysis, the pentose phosphate pathway, and the tryptophan/kynurenine pathway, without evidence of intestinal inflammation. Microbiome analysis revealed an increased abundance of the genus *Lactobacillus* in MS NASH mice. Colonic organoids from MetS mice showed altered goblet cell differentiation. Comparative analysis with human MetS fecal metabolomics demonstrated similar dysregulated pathways, underscoring the translational relevance of these findings. Our study reveals significant metabolic and microbial alterations in the colon of MS NASH mice, implicating a dysfunctional GI tract as a potential etiological factor in MetS. These findings highlight specific metabolic pathways and microbial signatures that could serve as future therapeutic targets for MetS.

**NEW & NOTEWORTHY:** This study identifies the colon as a metabolically active tissue affected in metabolic syndrome. Despite the absence of intestinal inflammation, MS NASH mice displayed altered colonic metabolism and microbiota composition, with conserved metabolite changes matching those seen in humans with metabolic syndrome. These findings highlight colonic metabolic dysfunction as a potential driver of gut dysbiosis and disease progression in metabolic syndrome and MASLD.

**Graphical Abstract:** 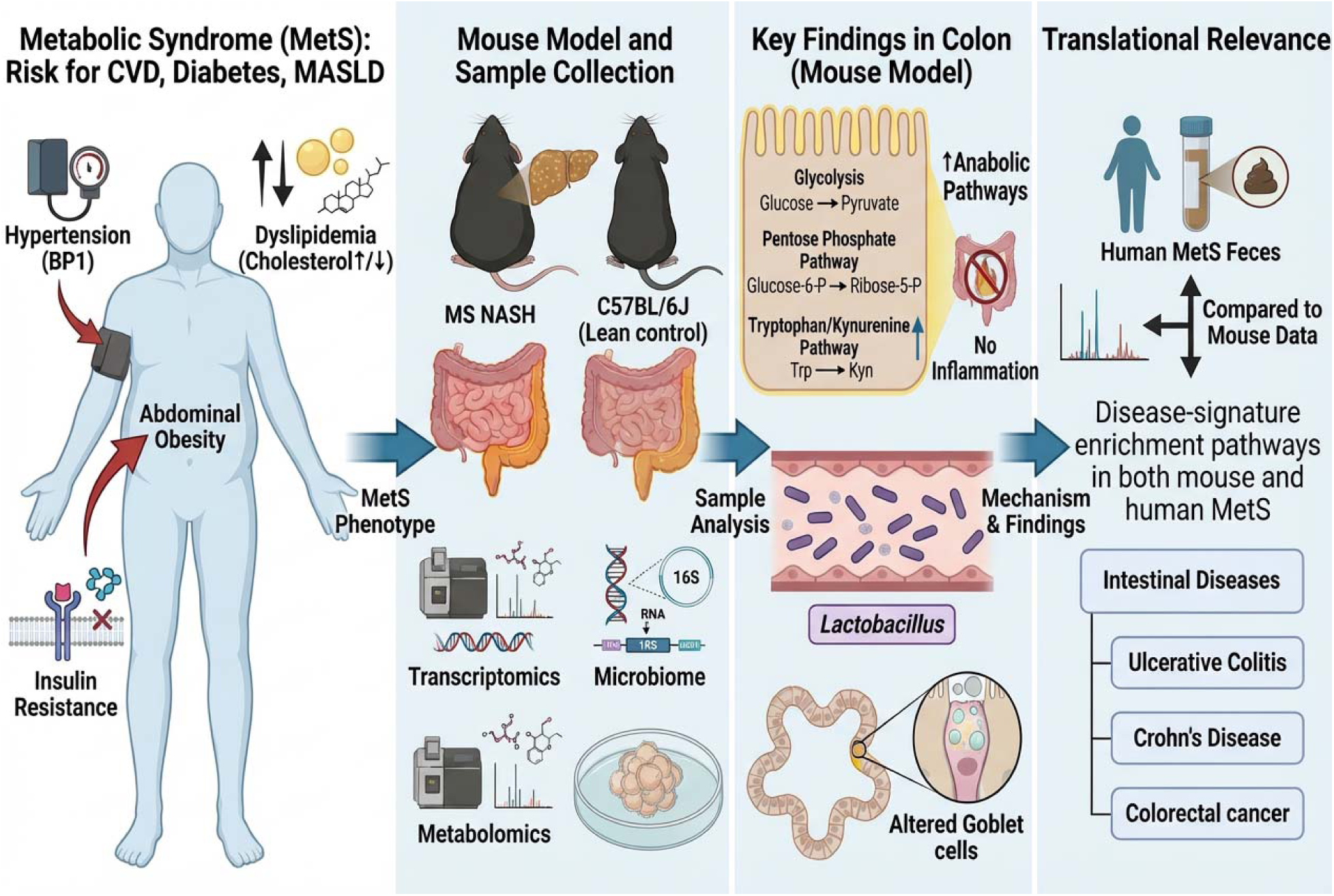

## INTRODUCTION

Metabolic Syndrome (MetS), a non-communicable disease, affects approximately one third of U.S. adults and an estimated quarter of the global population (1). MetS is diagnosed by the presence of at least three of the five risk factors: abdominal obesity, insulin resistance, hypertension, hypertriglyceridemia, and low high-density lipoprotein (HDL) cholesterol. Individuals with MetS have an increased risk to develop heart disease, type 2 diabetes (T2D), stroke, and metabolic dysfunction-associated steatotic liver disease (MASLD), formerly known as nonalcoholic fatty liver disease (2). Notably, MASLD is widely considered the hepatic manifestation of MetS (3, 4).

We recently reported individuals with MetS exhibit alterations in the fecal metabolome in the absence of overt intestinal inflammation, as assessed by fecal calprotectin, compared with non-MetS participants (5). Whether these changes occur at the cellular level remains unclear. In this brief report, we assessed the metabolic profile of the colon of MS NASH/PcoJ (herein called MS NASH) mice, a mouse model that has been used as a model of MetS (6–10). We report MS NASH mice show major alterations in the colon metabolome as determined by targeted transcriptomics and metabolomics. MS NASH mice also exhibited a distinct gut microbiota composition, characterized by clear community separation and disease-associated shifts in key bacterial genera. Lastly, integrated metabolite-set enrichment and cross-species analyses identified conserved alterations in energy- and mitochondria-associated metabolites that align MS NASH mouse colonic profiles with human MetS signatures. These findings demonstrate conserved metabolic alterations in the MS NASH colon that parallel human MetS signatures supporting the translational relevance of the model.

## MATERIALS AND METHODS

### Animals

The MS NASH/PcoJ (030888) (6–10) and C57BL/6J (000664) mice were purchased from Jackson Laboratory. All mice were 14-16 weeks of age and were male mice. All experiments were approved by the Institutional Animal Care and Use Committee of the University of New Mexico Health Sciences Center, in accordance with the National Institutes of Health guidelines for use of live animals. The University of New Mexico Health Sciences Center is accredited by the American Association for Accreditation of Laboratory Animal Care.

### Cells and Tissue Preparation

Upon necropsy, tissues were weighed, measured, sectioned out for various procedures, and flash frozen. Samples collected for downstream analysis included colon, liver, perigonadal fat pads, and stool. These samples were either flash frozen in liquid nitrogen and stored in −80C, fixed with formalin, or put into 70% ethanol (EtOH). Stool samples collected at euthanasia were processed and normalized to 90mg/mL using PBS + 0.1% TWEEN to analyze inflammatory markers of the colon such as Lipocalin-2 (LCN-2)(11) using R&D Systems Quantikine ELISA Mouse Lipocalin-2/NGAL Immunoassay (Catalog no. MLCN20). A section of the liver was stored in 10% formalin for 48 hours, then washed and stored in 70% EtOH prior to processing for tissue sectioning. Histopathological evaluation was performed on mouse liver tissue. Formalin-fixed, paraffin-embedded (FFPE) sections of livers were prepared at the University of New Mexico Comprehensive Cancer Center (UNMCCC) Human Tissue Repository (HTR) Core Facility. Sections (5 μm) were cut from blocks fixed in 10% neutral-buffered formalin and stained with hematoxylin and eosin (H&E) using standard protocols. For the generation of intestinal organoids, intestinal organoid media was comprised of Advanced Dulbecco’s modified Eagle medium/Ham’s F-12 (ThermoFisher, Waltham, MA), 100 U/mL penicillin/streptomycin (Quality Biological, Gaithersburg, MD), WNT surrogate-Fc fusion protein (ImmunoPrecise Antibodies Ltd, Utrecht, The Netherlands) 15% v/v R-spondin1 conditioned medium (cell line kindly provided by Calvin Kuo, Stanford University), 10% v/v Noggin conditioned medium (cell line kindly provided by Gijs van den Brink, Tytgat Institute for Liver and Intestinal Research), 1X B27 supplement (ThermoFisher), 10 mM HEPES (ThermoFisher), 1X GlutaMAX (ThermoFisher), 1mM N-acetylcysteine (MilliporeSigma), 50 ng/mL human epidermal growth factor (ThermoFisher), 10 nM [Leu-15] gastrin (AnaSpec, Fremont, CA), 500 nM A83-01 (Tocris, Bristol, United Kingdom), 10 μM SB202190 (MilliporeSigma), 100 mg/mL primocin (InvivoGen, San Diego, CA). Base media, for differentiation of organoids, had the same composition of media but lacked WNT surrogate Fc fusion protein, Rspo-1 and SB202190. Colonic crypt isolation and colonic organoid generation were prepared from proximal colons as previously reported (12–15). Isolated crypts were resuspended in Matrigel (Corning) and 25 µL droplets were plated in a 24-well tissue culture plate (Corning). After polymerization at 37°C, 500µL of organoid expansion media was added for 2 d. After 2 d, the organoid expansion media was replaced every other day. Colonic organoids were passaged every 3-7 d by harvesting in Cultrex Organoid Harvesting Solution (Bio-Techne, Minneapolis, MN) at 4°C with shaking for 45 min, as previously described (14, 15). All colonic organoids cultures were maintained at 37°C and 5% CO2. Unless noted, colonoid lines have been passaged >10 times. Colonic organoids were harvested from Matrigel using Cultrex.

### Microbiota analysis

Fresh fecal samples from C57BL/6J and MS NASH mice were collected fresh in sterile tubes and flash frozen. Microbial communities were determined by sequencing of the 16S rRNA as previously reported (11, 16) with minor modifications described below.

Microbial DNA was isolated from feces using the ZymoBIOMICS DNA Miniprep Kit (Zymo Research) following manufacturer’s recommendations. Variable regions V-3 through V-4 of the 16S rRNA gene were amplified by PCR using 100 ng input of DNA for each sample in duplicate using primers 319F- (5’-ACTCCTRCGGGAGGCAGCAG-3’) and 806R- (5’-GACAGGACTACHVGGGTATCTAATCC-3’) containing Nextera adapter overhangs. A second PCR was performed with Nextera® XT Index Kit v2 Set A (Illumina) to complete the adapter and add a unique sample-specific barcodes. After each PCR, a clean-up with AxyPrep Fragment Select-I magnetic beads (Axygen Biosciences) was completed, and all PCR reactions were run on an Applied Biosystems 2720 Thermal Cycler. Indexed samples were combined to yield duplicate 300 ng pools, followed by the creation and denaturation of a 4 nM library, and paired 250bp sequencing runs were completed on the Illumina MiSeq using v3 sequencing chemistry (Illumina). All reagents and kits are listed in Table S1.

### NanoString nCounter Gene Expression Analysis

Gene expression profiling was performed on proximal colon tissue using the NanoString nCounter XT platform (NanoString Technologies, Seattle, WA). Total RNA was isolated from proximal colon samples (n=12) and subjected to quality control prior to analysis. RNA quantity was initially assessed by NanoDrop spectrophotometry. Fluorometric quantification was performed using the Qubit RNA High Sensitivity assay (Thermo Fisher Scientific). RNA integrity and fragmentation profiles were evaluated using a Fragment Analyzer (Agilent Technologies). RNA QC summary metrics included total concentration, fragment distribution profiles, and assessment of 18S and 28S ribosomal RNA peaks. Because samples exhibited moderate RNA fragmentation, input RNA amounts were adjusted above the standard 100 ng recommended for intact RNA. Based on QC recommendations, 150-300 ng of total RNA per sample (typically 200-250 ng depending on degradation level) was used for hybridization. The same RNA input amount for each sample was maintained across both NanoString panels. NanoString Hybridization and Data Acquisition Gene expression profiling was performed at the University of Arizona Genetics Core using the NanoString nCounter XT platform (NanoString Technologies, Seattle, WA). Gene expression was assessed using a pre-designed NanoString mouse CodeSet panel: XT_Mm_Metabolic Pathways_CSO (Catalog no. 115000374). Reactions were performed using the nCounter XT Master Kit for 12 reactions (Catalog no. 100052). Hybridization was conducted at 65°C for 17 hours (overnight) according to manufacturer instructions. Following hybridization, samples were processed on the NanoString nCounter Prep Station and analyzed on the nCounter Digital Analyzer to obtain raw reporter code counts (RCC files). Reporter Library Files (RLFs) corresponding to each panel were used to map barcode counts to target transcripts during downstream analysis. For data processing and quality control, raw RCC files were imported into NanoString nSolver Analysis Software (NanoString Technologies). All samples passed initial NanoString quality control metrics, including imaging quality, binding density, positive control linearity, and limit of detection. Data normalization was performed using standard NanoString workflows, including: a) Positive control normalization to adjust for system variation and b) CodeSet content normalization using housekeeping genes. Normalized counts were used for downstream differential expression and pathway analyses.

### Metabolomics

Metabolomic profiling of colon tissue from C57BL/6J and MS NASH mice was performed by Human Metabolome Technologies, Inc. (HMT; Tsuruoka, Japan) using capillary electrophoresis–time-of-flight mass spectrometry (CE-TOFMS) (Table S1). Eight colon samples (n = 4 C57BL/6J, n = 4 MS NASH) were homogenized in 50% acetonitrile (v/v) containing 20 µM internal standards, centrifuged, and filtered through 5-kDa cut-off membranes to remove macromolecules. The filtrates were concentrated and resuspended in ultrapure water prior to analysis. Both cationic and anionic metabolites were analyzed under positive and negative electrospray ionization modes, respectively, using Agilent CE-TOFMS instruments (Agilent Technologies, Palo Alto, CA) equipped with fused-silica capillaries (50 µm × 80 cm). Analytical parameters included a capillary voltage of 30 kV, detector voltage of 4.0 kV, and a mass scan range of m/z 50–1000. Data were processed using MasterHands software (version 2.17.11; Keio University) for automated peak extraction, alignment, and normalization to internal standards. Metabolites were annotated by migration time (±0.5 min) and mass accuracy (±10 ppm) against HMT’s standard library. Relative peak areas were log_₂_-transformed and mean-centered, and metabolites detected in fewer than 75% of samples were excluded. Statistical analyses were performed using unpaired two-tailed Student’s t-tests (p < 0.05), and multivariate analysis by principal component analysis (PCA) and hierarchical clustering (Ward’s linkage, Euclidean distance) was used to visualize global metabolic differences between groups. Human stool metabolomics from individuals with and without metabolic syndrome was conducted by the NYU Langone Metabolomics Core Resource Laboratory (New York, NY) using a hybrid liquid chromatography–mass spectrometry (LC–MS) platform (Table S2). Approximately 10 mg of stool per subject was extracted in aqueous-organic solvent containing internal standards and analyzed on an Agilent 6545 Q-TOF mass spectrometer operating in both ESI+ and ESI− modes. Mass accuracy was maintained within ±1.2 ppm and retention-time stability within 0.1 min across all runs. Internal quality-control samples were used to monitor instrument precision and reproducibility. Data acquisition and peak processing were performed using Agilent MassHunter Qualitative Analysis and Profinder Suite (version 10.0). Metabolite identities were confirmed by accurate mass, isotopic distribution, and retention-time matching to NYU’s in-house MS/MS spectral library. Processed data were log_₂_-transformed, normalized to internal standards, and analyzed by unpaired two-tailed Student’s t-tests (p < 0.05). PCA and hierarchical clustering (Ward’s linkage, Euclidean distance) were used to assess group separation between healthy and metabolic-syndrome samples. For both mouse and human datasets, we used MetaboAnalyst v6.0 to run pathway enrichment analysis and check disease-signature enrichment using the feces metabolite-set library (Table S3). Enrichment statistics followed the globaltest method. Enrichment testing was conducted using the well-established globaltest algorithm, which calculates a Q-statistic for each metabolite set, incorporating generalized linear modeling to account for pathway topology. To directly compare directionality across species, we plotted log_₂_ fold-change (mouse: MS NASH vs C57BL/6J on the x-axis; human: MetS vs Control on the y-axis) for metabolites shared between datasets; per-metabolite p values were obtained with unpaired two-tailed t-tests and adjusted by Benjamini–Hochberg FDR, and metabolites were classified as increased or decreased when |log_₂_FC| ≥ 0.10 in both species. Only metabolite sets with five or more members were evaluated.

### RNA isolation, quantification, and RT-qPCR

RNA isolation was performed using the RNA Purelink Minikit (Invitrogen) according to the manufacturer’s protocols on colonic organoids which were harvested from Matrigel using Cultrex Organoid Harvesting Solution as previously described [5]. RNA was quantified on Nanodrop2000 and all samples yielded a 260/280 of 2 ± 0.15. cDNA synthesis was performed using SuperScript IV VILO Master Mix (ThermoFisher). Reverse transcription reaction utilized the viiA7 Thermal Cycler (ThermoFisher and RT-qPCR runs utilized Taqman MasterMix (ThermoFisher) using the QuantStudio 3. RT-qPCR primers (TaqMan Assays) and reagents used are listed in Table S4.

### Microscopy and image analysis

For confocal microscopy, macrophages were plated at 100k cells per well on 18 mm glass coverslips and stimulated with LPS (500 ng/mL) and IFN_ (10 ng/mL) for 8 hours and treated with BrefA (10 nM) or Baf. A1 (10 nM) for 2 hours. Cells were then fixed with 4% PFA followed by a wash with 1x PBS. Blocking buffer contained PBS with 50% FBS, 2% BSA and 0.1% saponin. After a 1-hour stain in primary antibody, cells were washed with PBS then followed by 1- hour stain in blocking buffer containing secondary antibody. Cells were mounted using ProLong Gold Antifade with DAPI and imaged using the Zeiss LSM 800 Airyscan Confocal microscope with a 63x oil objective lens. Images were processed using Zen Software and Adobe Photoshop (version CC 2019). All primary and secondary antibodies used for confocal staining are listed in Table S6.

### Statistical Analysis

Statistical analysis was performed as described in figure legends and graphs generated display mean (± SD) and were obtained using Prism software. Microbiome data was sequenced and processed by Illumina’s service lab using their in-house analysis pipeline. Cluster analysis was performed using heatmap3 (17) package in R. T-test was used to measure specific microbiome species abundance between conditions. Adjust p-value > 0.05 was used as significant threshold. Principle coordinate analysis was conducted in R. Confocal Images Statistical Analysis and Additional Software - Pearson’s Correlation Coefficient was acquired from BMM images using Huygens’s Deconvolution Scientific Volume Image Software (UNM Fluorescence Microscopy and Cell Imaging shared resource). Quantification figures were also made using Prism, while confocal image figures were constructed using Adobe Illustrator (version CC 2019). All other data were analyzed using one -way ANOVA or two-tailed unpaired Student’s t test (Prism).

## RESULTS

### MS NASH mice show phenotypic and microbial alterations in absence of intestinal inflammation

MS NASH mice are a polygenic mouse model of obesity, metabolic syndrome, MASLD and diabetes with an intact leptin pathway that have proven to be a translationally-relevant model (6–10, 18). MS NASH mice progressively develop several features of metabolic disorders that are similar to humans including steatosis, obesity, dyslipidemia, and insulin resistance (6–10, 18). Therefore, we examined both MS NASH and control (C57BL/6J; B6) mice after being on a standard diet for 10-weeks. MS NASH mice were significantly heavier than controls and exhibited increased perigonadal fat mass (**Fig. 1A-B**). MS NASH mice also displayed hepatomegaly (**Fig. 1C, H-I)** with histological evidence of macrovesicular steatosis **(Fig. 1F-G**), consistent with MASLD progression. Interestingly, MS NASH mice had significantly elongated colons compared to controls (**Fig. 1D**). Despite these pronounced metabolic and intestinal phenotypes, fecal lipocalin-2 (LCN-2) levels were unchanged between groups (**Fig. 1E**), indicating the absence of overt intestinal inflammation.

**Figure 1.**
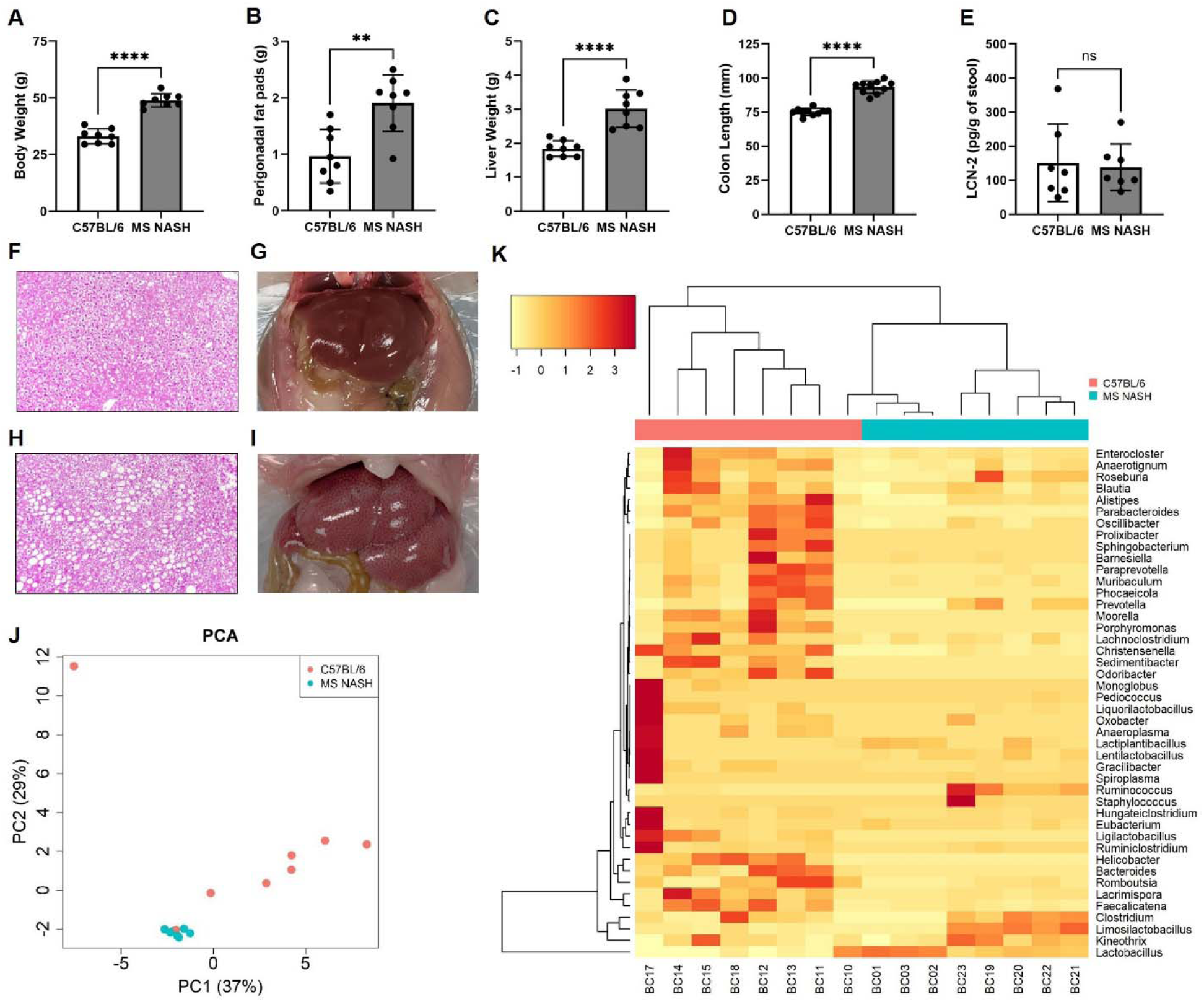
MS NASH mice exhibit metabolic and microbiota alterations without evidence of intestinal inflammation. (A-E) Metabolic and intestinal phenotypes in C57BL/6J (B6) control and MS NASH mice after 10 weeks on standard chow. (A) Body weight, (B) perigonadal fat pad weight, (C) liver weight, (D) colon length, and (E) fecal lipocalin-2 (LCN-2) concentrations. MS NASH mice exhibit increased body weight, adiposity, hepatomegaly, and colon length, while fecal LCN-2 levels are unchanged between groups. (F-I) Representative H&E-stained liver sections and gross liver images. Normal hepatic morphology in B6 controls (F) and macrovesicular steatosis in MS NASH mice (H). Representative gross liver images from B6 controls (G) and MS NASH mice (I), demonstrating hepatomegaly in MS NASH animals. (J) Principal coordinate analysis (PCA) of fecal microbial communities based on 16S rRNA sequencing showing separation between B6 and MS NASH mice (PC1 = 37%, PC2 = 29%). (K) Genus-level heatmap with hierarchical clustering demonstrating distinct microbial composition between groups. Data are presented as mean ± SEM with individual animals shown. Statistical significance was determined by unpaired two-tailed *t*-test. N = 8 mice per group.

To determine whether metabolic dysfunction was associated with alterations in the gut microbiota, we performed 16S rRNA sequencing on fecal samples from MS NASH and control mice. Principal coordinate analysis (PCoA) revealed clear separation between microbial communities, with PC1 and PC2 explaining 37% and 29% of the variance, respectively (**Fig. 1J**). Hierarchical clustering and genus-level heatmap analysis further demonstrated distinct microbial signatures between groups (**Fig. 1K**). Several taxa were differentially abundant between groups. Notably, members of the genus *Lactobacillus* and *Limosiolactobacillus* were increased in MS NASH mice, while multiple taxa including *Blautia*, *Parabacteroides*, *Prevotella*, and *Roseburia* were relatively enriched in controls. These alterations suggest a shift in the colonic microbiota composition in the context of metabolic syndrome, characterized by an expansion of lactobacilli and reduction of certain commensal anaerobes. Together, the PCA and hierarchical clustering results demonstrate that MS NASH mice exhibit significant gut microbiota remodeling, which may contribute to the metabolic and colonic alterations observed in this model.

### Colonic gene expression analysis reveals metabolic pathways reprogramming in MS NASH mice

To examine alterations in genes involved in core metabolic processes and signaling pathways, proximal colon tissues from B6 and MS NASH mice were profiled using the NanoString nCounter Metabolic Pathways gene expression panel. Genes were defined as upregulated or downregulated based on a ≥2-fold change in expression with a nominal P < 0.1 compared to control mice. Using these criteria, 298 genes were upregulated and 131 genes were downregulated in MS NASH mice (**Fig. 2 A-E**). Unsupervised hierarchical clustering of all 768 panel genes revealed clear segregation of MS NASH mice from controls, indicating widespread transcriptional remodeling of colonic metabolic programs (**Fig. 2F**). Pathway-level aggregation demonstrated prominent upregulation of carbon metabolism pathways, including glycolysis and the pentose phosphate pathway, as well as increased activity of the tryptophan-kynurenine metabolic axis (**Fig. 2G**). Complementary pathway enrichment analysis identified significant overrepresentation of gene sets associated with non-alcoholic fatty liver disease (now known as MASLD), oxidative phosphorylation, mitochondrial respiration, and carbon metabolism (**Fig. 2H**), consistent with enhanced bioenergetic demand and metabolic stress in the MS NASH colon. Collectively, these data demonstrate that the colonic epithelium of MS NASH mice undergoes broad changes in metabolic gene expression within the colon consistent with increased epithelia metabolic activity and metabolic stress.

**Figure 2.**
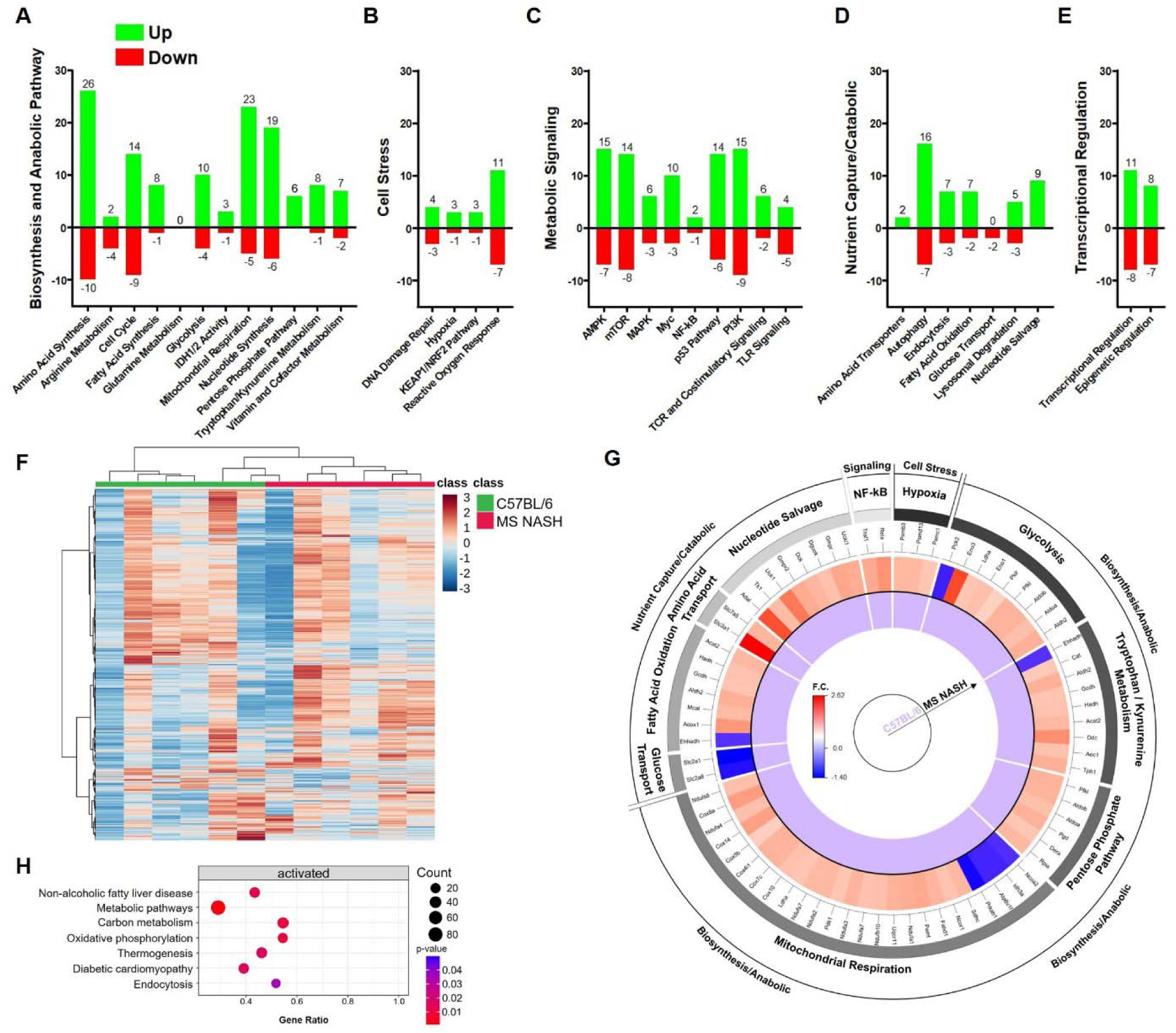
Colonic transcriptomic profiling reveals metabolic pathway reprogramming in MS NASH mice. (A-E) Number of differentially expressed genes in MS NASH mice relative to B6 controls across major metabolic themes: (A) biosynthesis/anabolic pathways, (B) cell stress pathways, (C) metabolic signaling pathways, (D) nutrient capture/catabolic pathways, and (E) transcriptional regulation. Green bars indicate upregulated genes and red bars indicate downregulated genes. Differential expression was defined as ≥2-fold change with nominal *P* < 0.1. (F) Heatmap of all 768 genes included in the NanoString nCounter Metabolic Pathways panel showing unsupervised hierarchical clustering of samples. Rows represent genes and columns represent individual mice. Data were autoscaled by gene, and clustering was performed using Pearson distance with Ward linkage. (G) Circos plot summarizing pathway-level transcriptional changes in MS NASH colons relative to controls. The outer ring denotes major metabolic themes, followed by individual metabolic pathways and gene-level fold changes. Color intensity represents log_2_ fold change (MS NASH vs B6). (H) Pathway enrichment analysis of differentially expressed genes identifying significantly activated metabolic and disease-associated pathways. Dot size represents gene count and color indicates nominal *P* value. N = 6 mice per group.

### Colonic metabolomic profiling reveals conserved pathway alterations between MS NASH mice and human metabolic syndrome

To complement the transcriptional data, targeted metabolomics was performed on colon tissues from B6 and MS NASH mice to define disease-associated metabolic alterations. Unsupervised clustering of metabolite abundances revealed clear separation between control and MS NASH colons, indicating widespread metabolic reprogramming in the MS NASH colon (**Fig. 3A**). *MetaboAnalyst* pathway enrichment analysis using the RaMP-DB metabolite and lipid library identified significant alterations in glycolysis/gluconeogenesis, the pentose phosphate pathway, tryptophan metabolism, and sphingolipid metabolism-pathways central to cellular energy homeostasis, redox balance, and inflammatory signaling (**Fig. 3B**). To assess translational conservation, disease-signature enrichment analysis was performed using the *MetaboAnalyst* fecal metabolite set library. The top 25 enriched disease-associated metabolite sets in MS NASH mouse colons closely mirrored those observed in human metabolic syndrome fecal metabolomics datasets (5), including signatures linked to inflammatory bowel disease, colorectal cancer, and metabolic dysfunction (**Fig. 3C–D**). Enrichment testing was conducted using the globaltest algorithm (19), which applies generalized linear modeling to evaluate coordinated metabolite-set behavior. Direct cross-species comparison of shared metabolites revealed concordant directional changes between MS NASH mice and human metabolic syndrome samples (**Fig. 3E**). Several metabolites were increased in both species, including 3-hydroxybutyric acid, uric acid, succinic acid, guanidinoacetic acid, N-acetylglutamic acid, and xanthosine. Notably, multiple concordantly elevated metabolites such as 3-hydroxybutyric acid and uric acid are established markers of mitochondrial dysfunction, altered energy metabolism, and MetS. Collectively, these data demonstrate that MS NASH mice exhibit colonic metabolomic and pathway-level alterations that closely parallel human MetS, supporting the translational relevance of this model for interrogating gut metabolic dysfunction in MASLD.

**Figure 3.**
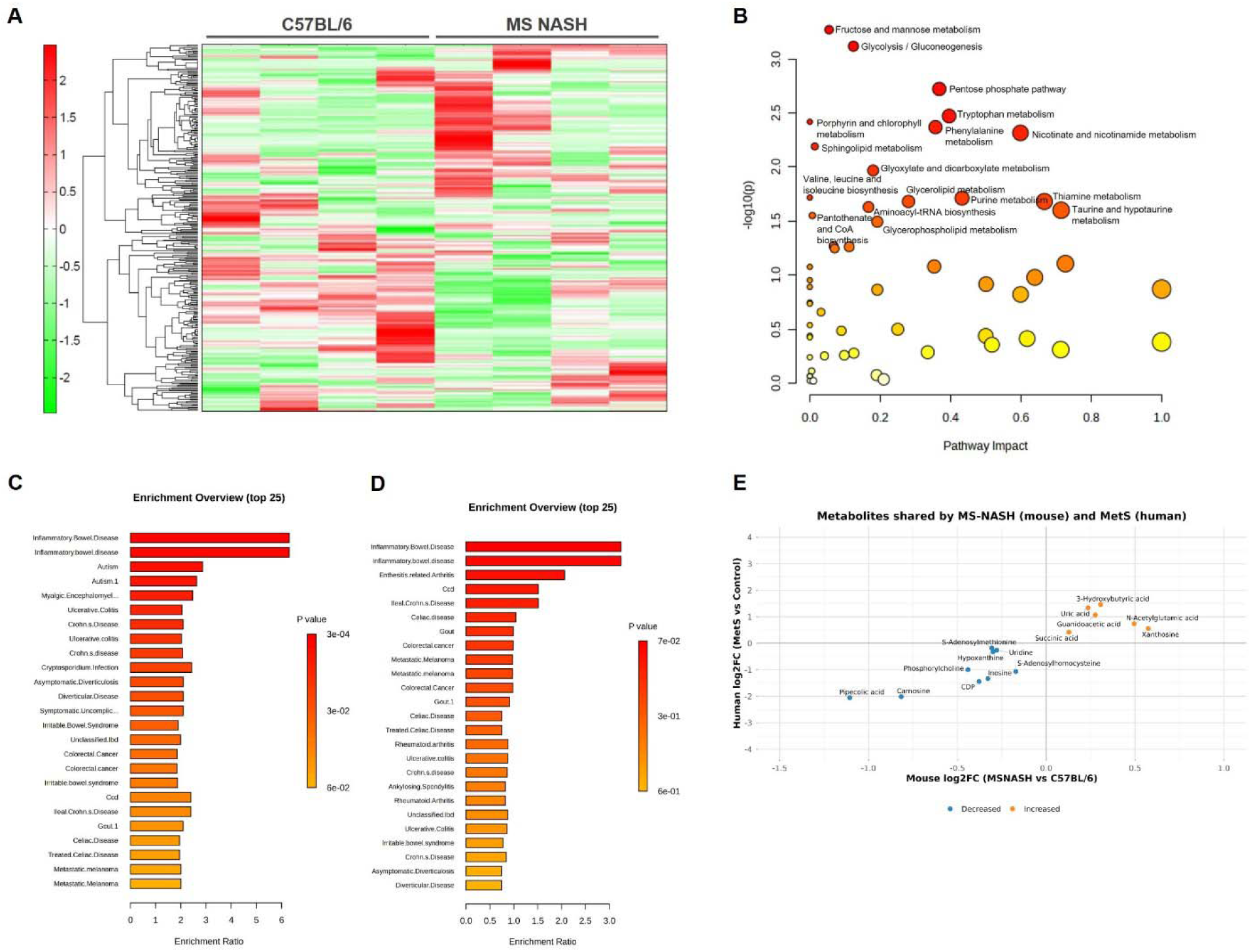
Colonic metabolomic alterations and conserved pathway enrichment between MS NASH mice and human metabolic syndrome. (A) Heatmap of colon metabolite abundances showing unsupervised hierarchical clustering of B6 (control) and MS NASH mice, demonstrating distinct metabolic profiles between groups. (B) Metabolic pathway impact analysis performed using MetaboAnalyst with the RaMP-DB metabolite and lipid library (integrating KEGG, HMDB, Reactome, and WikiPathways). Each point represents a metabolic pathway; the x-axis indicates pathway impact and the y-axis indicates -log10(*P*). Pathways with the greatest impact include glycolysis/gluconeogenesis, the pentose phosphate pathway, tryptophan metabolism, and sphingolipid metabolism. (C-D) *Disease-signature enrichment analysis* using the MetaboAnalyst fecal metabolite set library. (C) Top 25 enriched disease-associated metabolite sets derived from MS NASH mouse colon metabolomics. (D) Corresponding enrichment profile from human metabolic syndrome fecal metabolomics datasets. Enrichment testing was performed using the globaltest algorithm; only metabolite sets containing ≥5 metabolites were included. (E) Cross-species comparison of metabolites shared between MS NASH mouse colons and human metabolic syndrome stool datasets. Log_2_ fold change values were calculated from group means (MS NASH vs B6 on the x-axis; MetS vs control on the y-axis). Metabolites were classified as increased (orange) or decreased (blue) when |log_2_FC| ≥ 0.10 in both species following Benjamini-Hochberg false discovery rate correction of per-metabolite *P* values (unpaired two-tailed *t*-tests).

### Colonoids derived from MS NASH mice exhibit altered epithelial marker expression under stem cell and differentiated conditions

To determine whether metabolic disease is associated with intrinsic changes in colonic epithelial cell phenotype, we generated colonic organoids (colonoids) from B6 and MS NASH mice. Colonoids were maintained under intestinal stem cell (ISC) conditions or induced to differentiate by withdrawal of WNT3A and R-spondin (**Fig. 4A-E**). Gene expression analysis demonstrated genotype-specific differences in epithelial lineage markers. Under ISC conditions, transcripts encoding the ISC marker *Lgr5* (**Fig. 4A**) and the goblet cell marker *Muc2* (**Fig. 4D**) were differentially expressed between control and MS NASH colonoids. Upon differentiation following WNT3A and R-spondin withdrawal, *Lgr5* expression was significantly reduced in both control and MS NASH colonoids (**Fig. 4C-D**), confirming effective induction of differentiation. Under these conditions, expression of *Muc2* (**Fig. 4E**) significantly altered in MS NASH colonoids relative to controls. Consistent with these transcriptional differences, immunofluorescence analysis revealed MS NASH colonoids demonstrated markedly increased MUC2 signal compared to controls, suggesting enhanced goblet cell-associated features even under differentiation conditions (**Fig. 4F-G**). Collectively, these findings indicate that epithelial cells from MS NASH mice exhibit intrinsic alterations in lineage specification, characterized by a persistent bias toward secretory cell programs across stem cell and differentiated states. This intrinsic epithelial remodeling provides a mechanistic link between metabolic disease and altered intestinal epithelial function independent of inflammatory cues.

**Figure 4.**
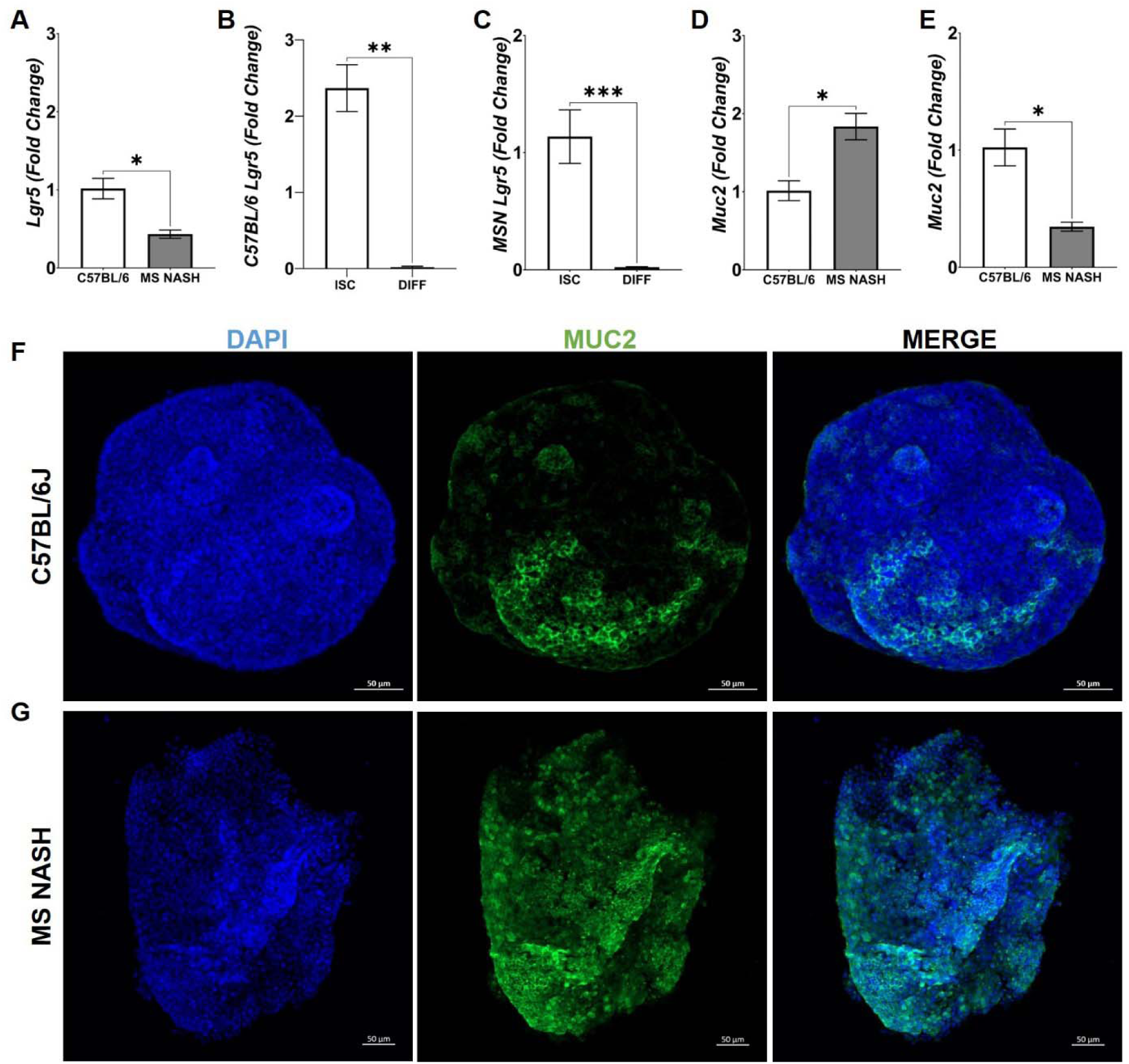
Colonoids derived from MS NASH mice exhibit altered epithelial lineage marker expression. (A-E) Gene expression of the intestinal stem cell marker *Lgr5* and the goblet cell marker *Muc2* in colonoids derived from B6 and MS NASH mice maintained under intestinal stem cell (ISC) culture conditions or following differentiation induced by withdrawal of WNT3A and R-spondin. (A) Comparison of *Lgr5* expression in ISC colonoids derived from B6 and MS NASH mice. (B-C) *Lgr5* expression in ISC versus differentiated colonoids in B6 (B) or MS NASH (C), demonstrating reduced stem cell marker expression following differentiation. (D) Comparison of *Muc2* expression in ISC colonoids derived from B6 and MS NASH mice. (E) Comparison of *Muc2* expression in differentiated colonoids derived from B6 and MS NASH mice. (F-G) Representative immunofluorescence images of colonoids stained for MUC2 demonstrating increased MUC2 signal in MS NASH colonoids compared with controls. Data are presented as mean ± SEM with individual biological replicates shown. N = 3 mice per group. Statistical significance was determined using an unpaired two-tailed *t*-test.

## DISCUSSION

In the present study, we demonstrate that the MS NASH mouse model develops pronounced metabolic and intestinal alterations in the absence of overt intestinal inflammation (6–10). Consistent with prior reports MS NASH mice displayed hallmark features of MetS including obesity, increased visceral adiposity, hepatomegaly, and hepatic steatosis. Despite these systemic metabolic abnormalities, fecal LCN-2 levels were unchanged similar to our previous findings with fecal calprotectin in individuals with MetS (5), suggesting that intestinal remodeling occurs independently of classical inflammatory responses. Instead, MS NASH mice exhibited significant restructuring of the gut microbiota, characterized by enrichment of *Lactobacillus* and *Limosiolactobacillus* and reduced abundance of several commensal anaerobes including *Blautia*, *Parabacteroides*, *Prevotella*, and *Roseburia*. At the molecular level, transcriptomic profiling revealed widespread alterations in metabolic gene expression within the colon, with increased expression of genes involved in carbon metabolism, mitochondrial respiration, and tryptophan-kynurenine metabolism. These transcriptional alterations were accompanied by concordant metabolomic changes including increased metabolites associated with mitochondrial stress and altered energy metabolism. Importantly, these metabolic signatures closely mirrored those reported in human MetS metabolomics dataset (5). Finally, colonoids derived from MS NASH mice exhibited persistent alterations in epithelial lineage marker expression including enhanced goblet cell-associated features that suggest metabolic disease induces intrinsic epithelial remodeling independent of systemic or immune-derived cues.

There is emerging evidence that MetS and MASLD are associated with profound alterations in the gut microbiota and intestinal metabolic function. Several studies have reported microbiota shifts in various metabolic diseases characterized by reductions in butyrate-producing taxa such as *Roseburia* and *Blautia* and an expansion of facultative anaerobes or lactobacilli (20–25). These changes have been linked to altered intestinal energy metabolism and host metabolic dysfunction (26–28). Consistent with these observations, the depletion of short chain fatty acid-producing bacteria observed in our study may contribute to epithelial metabolic stress and altered colonocyte bioenergetics. Our transcriptional and metabolic data further support the concept that the intestinal epithelium undergoes metabolic rewiring in metabolic diseases. Increased activity of glycolysis, the pentose phosphate pathway, and mitochondrial respiration has been described in metabolically stressed epithelial tissues, which may reflect adaptive responses to increased energetic demand or oxidative stress (29–34). Additionally, the observed activation of the tryptophan-kynurenine pathway is notable, as this pathway has been implicated in immune modulation, epithelial homeostasis, and metabolic disease progression (35–37). These findings indicate that MetS reshapes intestinal physiology through coordinated alterations in microbial composition and epithelial metabolic reprogramming.

An important observation in this study is that epithelial alterations persisted in colonoids derived from MS NASH mice, even after removal from the in vivo metabolic environment. This suggests that metabolic disease may induce stable epithelial reprogramming potentially through epigenetic remodeling in ISC. Similar findings have been suggested in other metabolic and inflammatory settings where ISC populations acquire persistent transcriptional programs that influences lineage differentiation and epithelial barrier function (38–40). Nevertheless, several limitations should be considered. First, the microbiota analysis relied on 16S rRNA sequencing, which limits taxonomic resolution and does not capture functional microbial activity. Second, the metabolomics profiling was targeted and may not capture the full spectrum of metabolic alterations present in the colon. While the colonoid experiments demonstrate intrinsic epithelial changes characterization of human colonoids derived from individuals with MetS are needed. Future studies integrating metagenomics, spatial metabolomics, and host-microbial co-culture systems could further define the mechanistic links between microbial metabolites, epithelial metabolism, and metabolic disease progression.

In summary, our work demonstrates that MetS and MASLD are associated with coordinated alterations in gut microbiota composition, epithelial metabolic pathways, and intestinal epithelial lineage programs in the absence of overt intestinal inflammation. The close alignment between metabolomic signatures observed in MS NASH mice and human MetS datasets underscores the translational relevance of this model for studying gut metabolic dysfunction. Our findings further suggest that metabolic disease can induce persistent epithelial reprogramming that may contribute to altered intestinal physiology and host-microbiome interactions. These findings highlight the intestine as an important metabolic organ in systemic metabolic disease and suggest that gut metabolic pathways or microbiota-derived metabolites may represent promising therapeutic strategies for MetS and MASLD.

## Acknowledgments

We further acknowledge Yan Guo and the UNM Comprehensive Cancer Center Support Grant NCI P30CA118100 and the HSC Shared Equipment for use of the Olympus iX83 Yokogawa Spinning Disk Confocal Microscope and Evident APEXVIEW APX100 Benchtop Fluorescence Microscope within the Advanced Light Microscopy Resource and the Bioinformatics Shared Resource at University of New Mexico Comprehensive Cancer Center with grant P30CA118100. We thank Michael L. Paffett at the UNM Advanced Light Microscopy Resource for training and guidance with confocal imaging. We thank Julie In for providing reagents for organoid experiments. We acknowledge the Human Metabolome Technologies, Inc and NYU Langone Health’s Metabolomics Laboratory for its help in acquiring the data presented. The data underlying this article are available in the article and in its online supplementary material.

## Author contributions

JAR, DPS, and EPH performed all analysis with the help from CME, SDM, ALN, RRG, and EFC. SDM and EFC contributed to microbiota sequencing and analysis. JAR and EFC participated in writing the manuscript. EFC designed the study, analyzed data and wrote the paper. All authors approved the final version of the manuscript.

## REFERENCES

1. Sherling DH, Perumareddi P, and Hennekens CH. Metabolic Syndrome. J Cardiovasc Pharmacol Ther 22: 365–367, 2017.

2. Kim D, Touros A, and Kim WR. Nonalcoholic Fatty Liver Disease and Metabolic Syndrome. Clin Liver Dis 22: 133–140, 2018.

3. Goyal A, Arora H, and Arora S. Prevalence of fatty liver in metabolic syndrome. J Family Med Prim Care 9: 3246–3250, 2020.

4. Yki-Jarvinen H. Non-alcoholic fatty liver disease as a cause and a consequence of metabolic syndrome. Lancet Diabetes Endocrinol 2: 901–910, 2014.

5. Coleman MJ, Espino LM, Lebensohn H, Zimkute MV, Yaghooti N, Ling CL, Gross JM, Listwan N, Cano S, Garcia V, Lovato DM, Tigert SL, Jones DR, Gullapalli RR, Rakov NE, Torrazza Perez EG, and Castillo EF. Individuals with Metabolic Syndrome Show Altered Fecal Lipidomic Profiles with No Signs of Intestinal Inflammation or Increased Intestinal Permeability. Metabolites 12: 2022.

6. Boland ML, Oldham S, Boland BB, Will S, Lapointe JM, Guionaud S, Rhodes CJ, and Trevaskis JL. Nonalcoholic steatohepatitis severity is defined by a failure in compensatory antioxidant capacity in the setting of mitochondrial dysfunction. World J Gastroenterol 24: 1748–1765, 2018.

7. Droz BA, Sneed BL, Jackson CV, Zimmerman KM, Michael MD, Emmerson PJ, Coskun T, and Peterson RG. Correlation of disease severity with body weight and high fat diet in the FATZO/Pco mouse. PLoS One 12: e0179808, 2017.

8. Peterson RG, Jackson CV, Zimmerman KM, Alsina-Fernandez J, Michael MD, Emmerson PJ, and Coskun T. Glucose dysregulation and response to common anti-diabetic agents in the FATZO/Pco mouse. PLoS One 12: e0179856, 2017.

9. Sun G, Jackson CV, Zimmerman K, Zhang LK, Finnearty CM, Sandusky GE, Zhang G, Peterson RG, and Wang YJ. The FATZO mouse, a next generation model of type 2 diabetes, develops NAFLD and NASH when fed a Western diet supplemented with fructose. BMC Gastroenterol 19: 41, 2019.

10. Zhang G, Wang X, Chung TY, Ye W, Hodge L, Zhang L, Chng K, Xiao YF, and Wang YJ. Carbon tetrachloride (CCl4) accelerated development of non-alcoholic fatty liver disease (NAFLD)/steatohepatitis (NASH) in MS-NASH mice fed western diet supplemented with fructose (WDF). BMC Gastroenterol 20: 339, 2020.

11. Merkley SD, Goodfellow SM, Guo Y, Wilton ZER, Byrum JR, Schwalm KC, Dinwiddie DL, Gullapalli RR, Deretic V, Jimenez Hernandez A, Bradfute SB, In JG, and Castillo EF. Non-autophagy Role of Atg5 and NBR1 in Unconventional Secretion of IL-12 Prevents Gut Dysbiosis and Inflammation. J Crohns Colitis 16: 259–274, 2022.

12. In JG, Yin J, Atanga R, Doucet M, Cole RN, DeVine L, Donowitz M, Zachos NC, Blutt SE, Estes MK, and Kovbasnjuk O. Epithelial WNT2B and Desert Hedgehog Are Necessary for Human Colonoid Regeneration after Bacterial Cytotoxin Injury. iScience 23: 101618, 2020.

13. Atanga R, Romero AS, Hernandez AJ, Peralta-Herrera E, Merkley SD, In JG, and Castillo EF. Inflammatory macrophages prevent colonic goblet and enteroendocrine cell differentiation through Notch signaling. bioRxiv 2023.

14. Atanga R, Parra AS, and In JG. Efficient RNA and RNA-protein co-detection in 3D colonoids by whole-mount staining. STAR Protoc 3: 101775, 2022.

15. Atanga R, Appell LL, Thompson MN, Lauer FT, Brearley A, Campen MJ, Castillo EF, and In JG. Single Cell Analysis of Human Colonoids Exposed to Uranium-Bearing Dust. Environ Health Perspect 132: 57006, 2024.

16. Komesu YM, Richter HE, Dinwiddie DL, Siddiqui NY, Sung VW, Lukacz ES, Ridgeway B, Arya LA, Zyczynski HM, Rogers RG, and Gantz M. Methodology for a vaginal and urinary microbiome study in women with mixed urinary incontinence. Int Urogynecol J 28: 711–720, 2017.

17. Zhao S, Guo Y, Sheng Q, and Shyr Y. Advanced heat map and clustering analysis using heatmap3. Biomed Res Int 2014: 986048, 2014.

18. Neff EP. Farewell, FATZO: a NASH mouse update. Lab Anim (NY) 48: 151, 2019.

19. Xu N, Solari A, and Goeman JJ. Closed testing with Globaltest, with application in metabolomics. Biometrics 79: 1103–1113, 2023.

20. Huang W, Zhu W, Lin Y, Chan FKL, Xu Z, and Ng SC. Roseburia hominis improves host metabolism in diet-induced obesity. Gut Microbes 17: 2467193, 2025.

21. Jiang S, Xie S, Lv D, Zhang Y, Deng J, Zeng L, and Chen Y. A reduction in the butyrate producing species Roseburia spp. and Faecalibacterium prausnitzii is associated with chronic kidney disease progression. Antonie Van Leeuwenhoek 109: 1389–1396, 2016.

22. Karlsson FH, Tremaroli V, Nookaew I, Bergstrom G, Behre CJ, Fagerberg B, Nielsen J, and Backhed F. Gut metagenome in European women with normal, impaired and diabetic glucose control. Nature 498: 99–103, 2013.

23. Drissi F, Raoult D, and Merhej V. Metabolic role of lactobacilli in weight modification in humans and animals. Microb Pathog 106: 182–194, 2017.

24. Niu Y, Hu X, Song Y, Wang C, Luo P, Ni S, Jiao F, Qiu J, Jiang W, Yang S, Chen J, Huang R, Jiang H, Chen S, Zhai Q, Xiao J, and Guo F. Blautia Coccoides is a Newly Identified Bacterium Increased by Leucine Deprivation and has a Novel Function in Improving Metabolic Disorders. Adv Sci (Weinh) 11: e2309255, 2024.

25. Hosomi K, Saito M, Park J, Murakami H, Shibata N, Ando M, Nagatake T, Konishi K, Ohno H, Tanisawa K, Mohsen A, Chen YA, Kawashima H, Natsume-Kitatani Y, Oka Y, Shimizu H, Furuta M, Tojima Y, Sawane K, Saika A, Kondo S, Yonejima Y, Takeyama H, Matsutani A, Mizuguchi K, Miyachi M, and Kunisawa J. Oral administration of Blautia wexlerae ameliorates obesity and type 2 diabetes via metabolic remodeling of the gut microbiota. Nat Commun 13: 4477, 2022.

26. Tilg H, and Moschen AR. Mechanisms behind the link between obesity and gastrointestinal cancers. Best Pract Res Clin Gastroenterol 28: 599–610, 2014.

27. Boursier J, Mueller O, Barret M, Machado M, Fizanne L, Araujo-Perez F, Guy CD, Seed PC, Rawls JF, David LA, Hunault G, Oberti F, Cales P, and Diehl AM. The severity of nonalcoholic fatty liver disease is associated with gut dysbiosis and shift in the metabolic function of the gut microbiota. Hepatology 63: 764–775, 2016.

28. Le Roy T, Llopis M, Lepage P, Bruneau A, Rabot S, Bevilacqua C, Martin P, Philippe C, Walker F, Bado A, Perlemuter G, Cassard-Doulcier AM, and Gerard P. Intestinal microbiota determines development of non-alcoholic fatty liver disease in mice. Gut 62: 1787–1794, 2013.

29. Litvak Y, Byndloss MX, and Baumler AJ. Colonocyte metabolism shapes the gut microbiota. Science 362: 2018.

30. Dias Nirello V, Araujo N, Carvalho de Assis H, Moreno-Gonzalez M, Ruiz P, Castro PR, Shealy NG, Shelton C, Font Fernandes M, de Oliveira S, Boroni M, Ryffel B, Byndloss MX, Beraza N, Ramirez Vinolo MA, and Varga-Weisz P. Microbiota shape the colon epithelium controlling inter-crypt absorptive goblet cells via butyrate-GP R109A signalling. Gut Microbes 17: 2573045, 2025.

31. Yoo W, Zieba JK, Foegeding NJ, Torres TP, Shelton CD, Shealy NG, Byndloss AJ, Cevallos SA, Gertz E, Tiffany CR, Thomas JD, Litvak Y, Nguyen H, Olsan EE, Bennett BJ, Rathmell JC, Major AS, Baumler AJ, and Byndloss MX. High-fat diet-induced colonocyte dysfunction escalates microbiota-derived trimethylamine N-oxide. Science 373: 813–818, 2021.

32. Kierans SJ, Fagundes RR, Malkov MI, Sparkes R, Dillon ET, Smolenski A, Faber KN, and Taylor CT. Hypoxia induces a glycolytic complex in intestinal epithelial cells independent of HIF-1-driven glycolytic gene expression. Proc Natl Acad Sci U S A 120: e2208117120, 2023.

33. Chiba K, Nomoto H, Izumihara R, Zhang X, Kameda H, Nakamura A, and Atsumi T. Metabolic adaptation to acute metabolic stress via PFKFB3 upregulation in rodent beta cells. Front Endocrinol (Lausanne) 16: 1552700, 2025.

34. TeSlaa T, Ralser M, Fan J, and Rabinowitz JD. The pentose phosphate pathway in health and disease. Nat Metab 5: 1275–1289, 2023.

35. Taleb S. Tryptophan Dietary Impacts Gut Barrier and Metabolic Diseases. Front Immunol 10: 2113, 2019.

36. Wang S, van Schooten FJ, Jin H, Jonkers D, and Godschalk R. The Involvement of Intestinal Tryptophan Metabolism in Inflammatory Bowel Disease Identified by a Meta-Analysis of the Transcriptome and a Systematic Review of the Metabolome. Nutrients 15: 2023.

37. Chajadine M, Laurans L, Radecke T, Mouttoulingam N, Al-Rifai R, Bacquer E, Delaroque C, Rytter H, Bredon M, Knosp C, Vilar J, Fontaine C, Suffee N, Vandestienne M, Esposito B, Dairou J, Launay JM, Callebert J, Tedgui A, Ait-Oufella H, Sokol H, Chassaing B, and Taleb S. Harnessing intestinal tryptophan catabolism to relieve atherosclerosis in mice. Nat Commun 15: 6390, 2024.

38. Singh V, Johnson K, Yin J, Lee S, Lin R, Yu H, In J, Foulke-Abel J, Zachos NC, Donowitz M, and Rong Y. Chronic Inflammation in Ulcerative Colitis Causes Long-Term Changes in Goblet Cell Function. Cell Mol Gastroenterol Hepatol 13: 219–232, 2022.

39. Nusse YM, Savage AK, Marangoni P, Rosendahl-Huber AKM, Landman TA, de Sauvage FJ, Locksley RM, and Klein OD. Parasitic helminths induce fetal-like reversion in the intestinal stem cell niche. Nature 559: 109–113, 2018.

40. Peck BC, Mah AT, Pitman WA, Ding S, Lund PK, and Sethupathy P. Functional Transcriptomics in Diverse Intestinal Epithelial Cell Types Reveals Robust MicroRNA Sensitivity in Intestinal Stem Cells to Microbial Status. J Biol Chem 292: 2586–2600, 2017.

